## Supplement Materials for "Deciphering Abnormal Platelet Subpopulations in Inflammatory Diseases through Machine Learning and Single-Cell Transcriptomics"

A

B

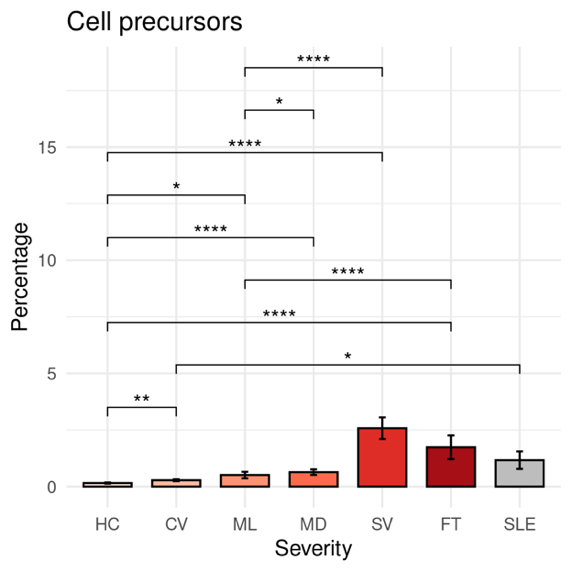

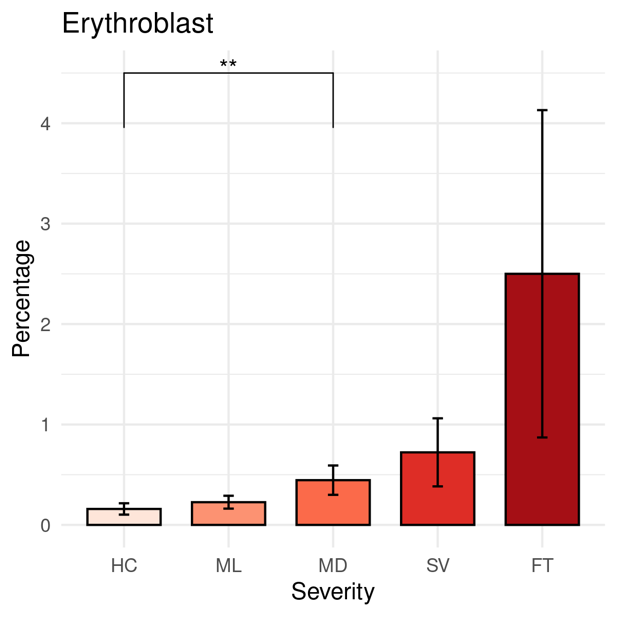

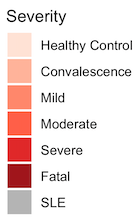

**Supplementary Figure 1**. **PBMC profiling from healthy controls, sepsis, similar symptom hospitalized, COVID-19 and SLE patients.** (A-C) Bar plots depicting the percentage of different cell types under different disease severities. A) Cell precursors, B) Erythroblast

The differences in percentages associated with adjusted *P*-values below 0.05, 0.01, 0.001, and 0.0001 are indicated as *, **, ***, and ****, respectively and not significant are not shown. The significance analysis was performed using Wilcoxon tests. Standard error bars were also added.

A

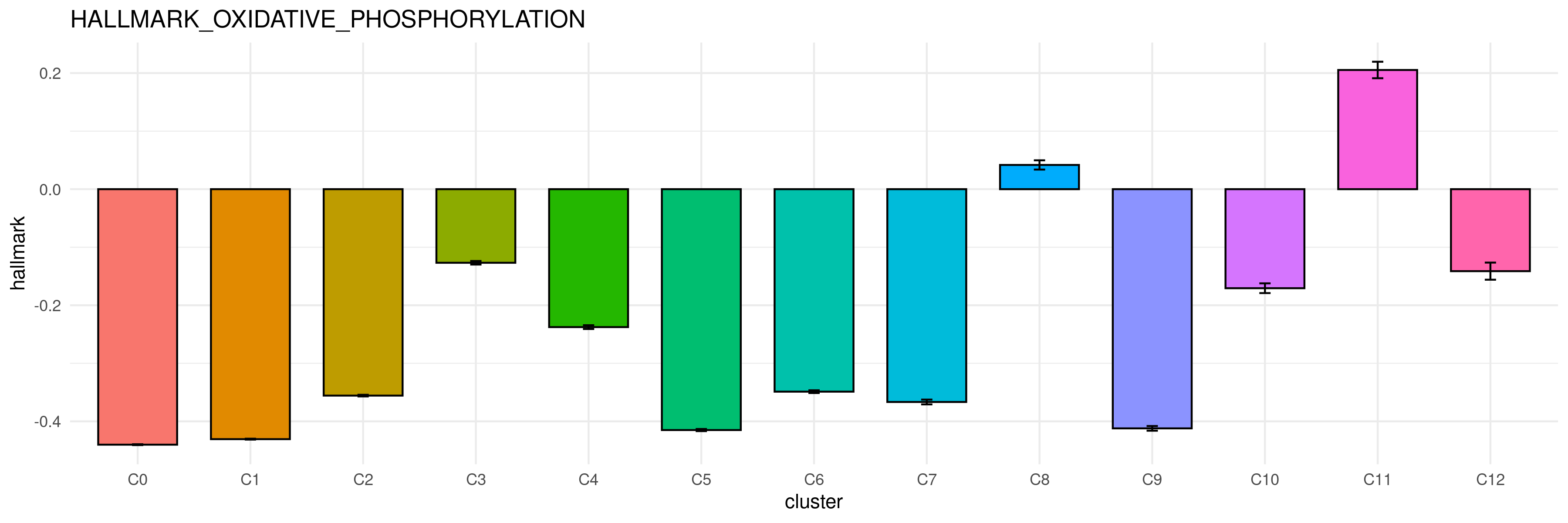

B

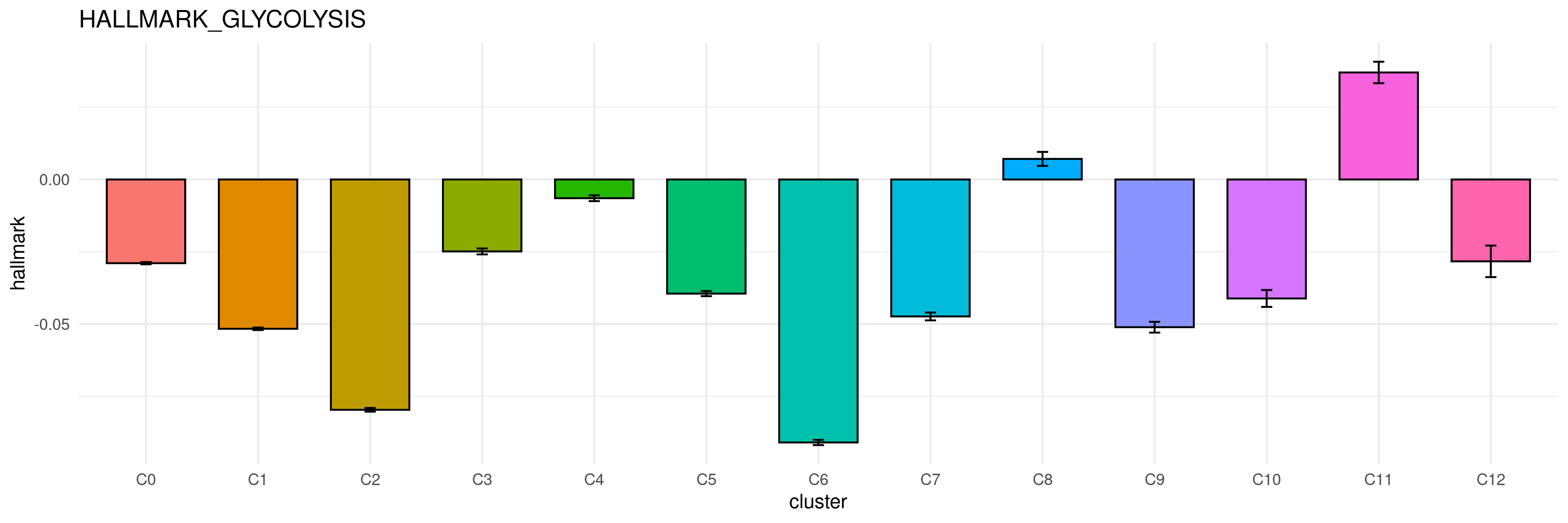

C

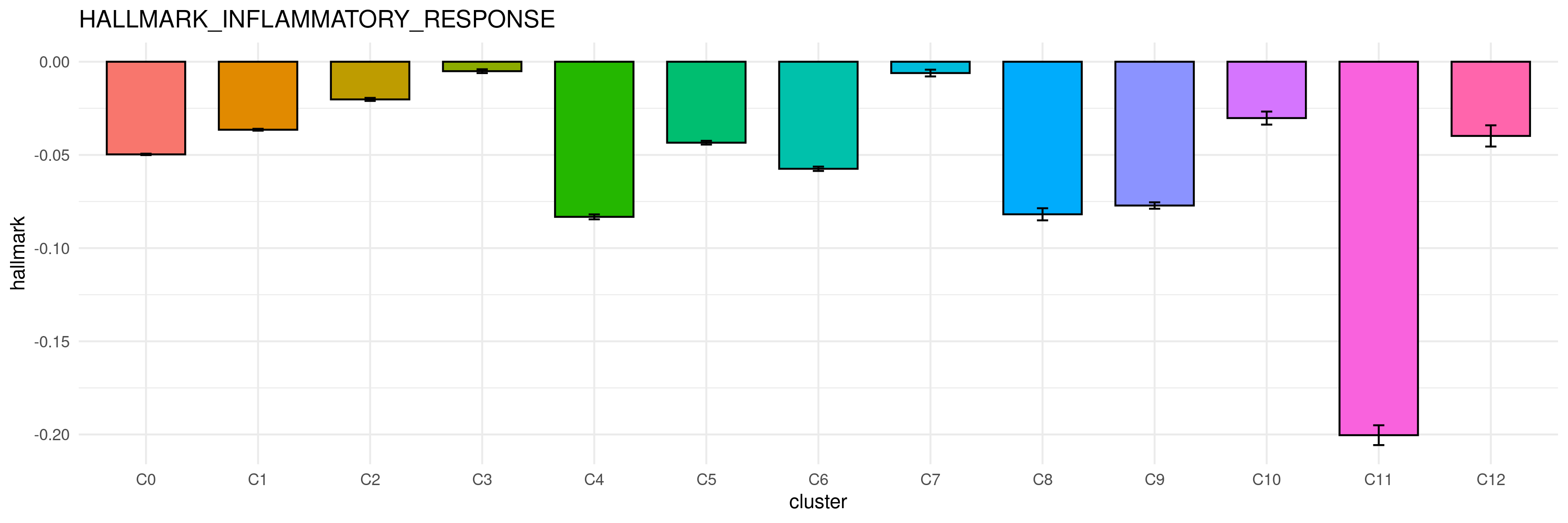

D

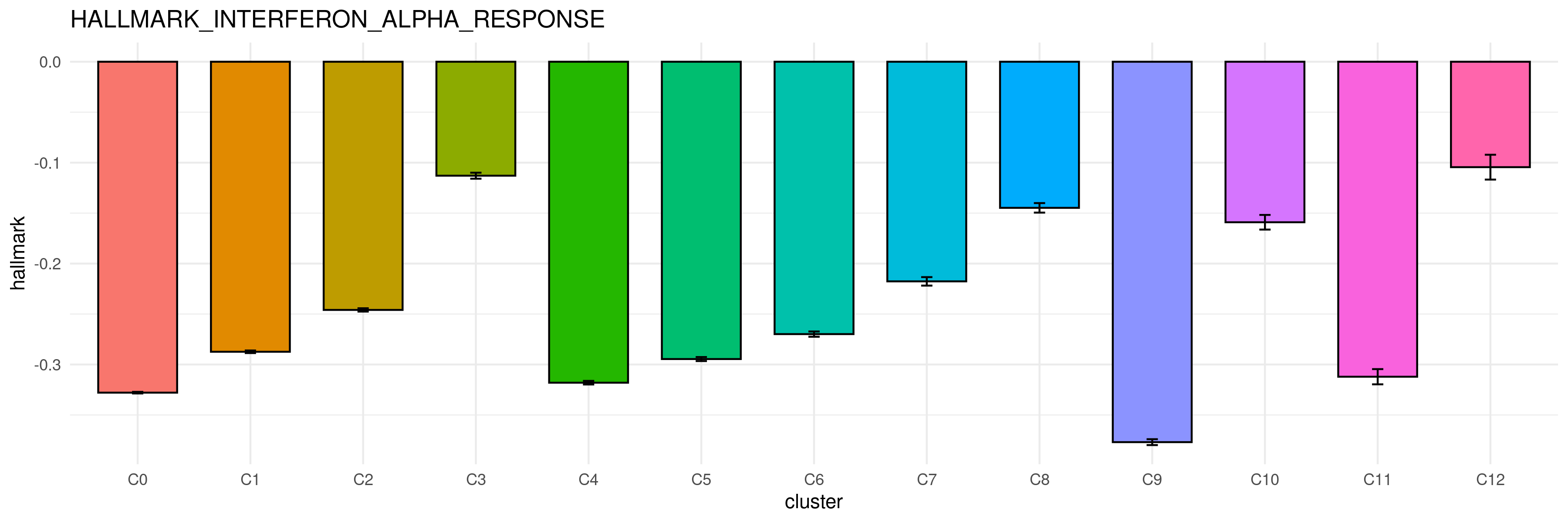

E

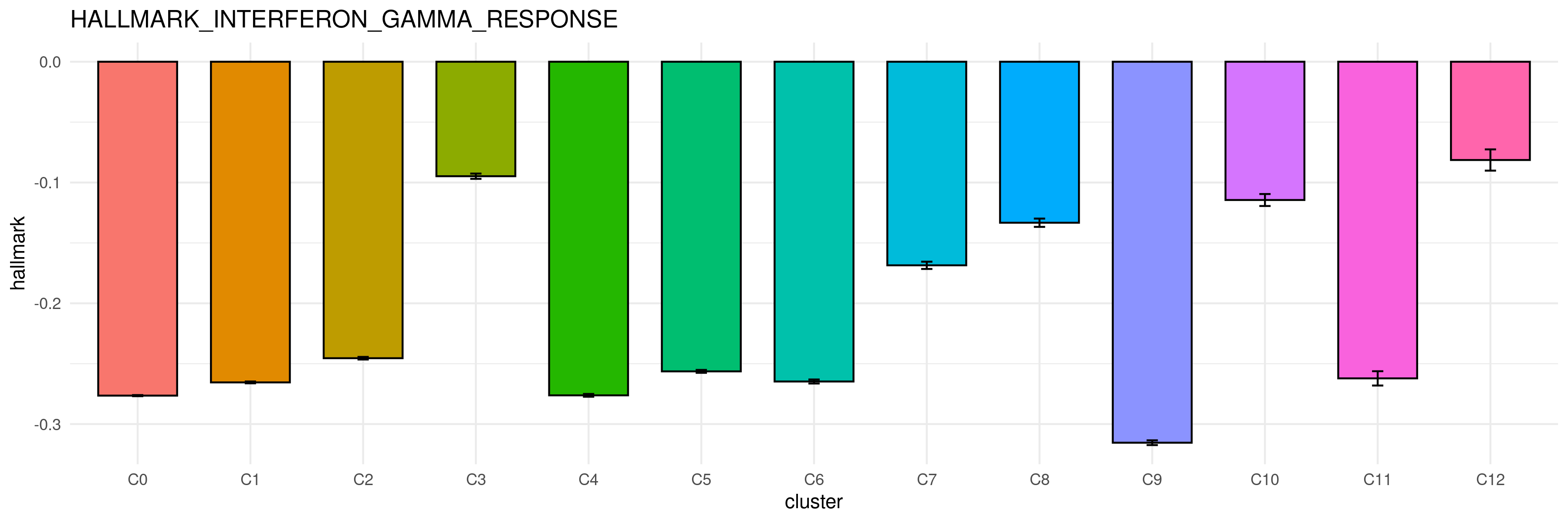

F

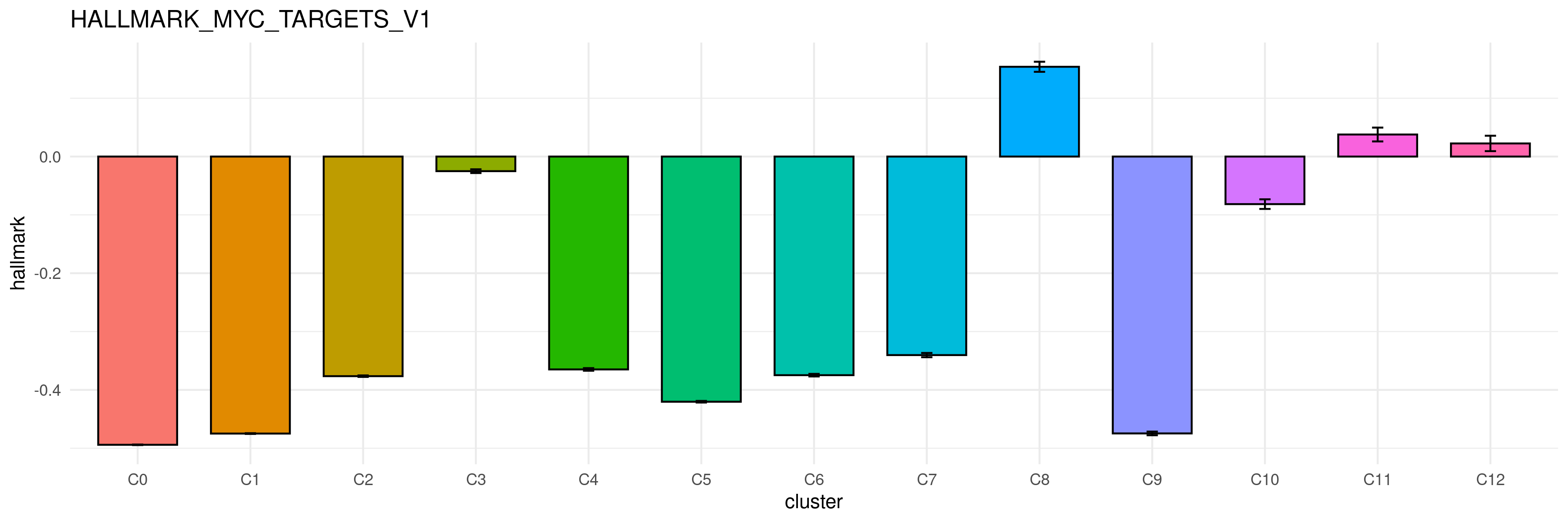

G

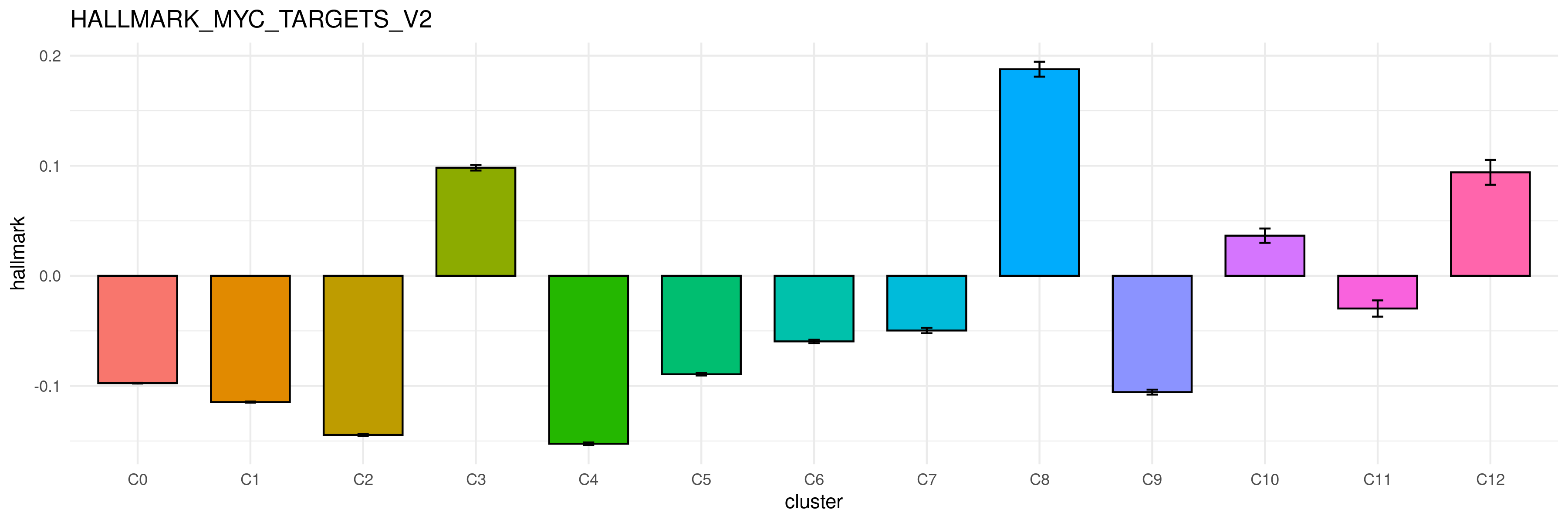

H

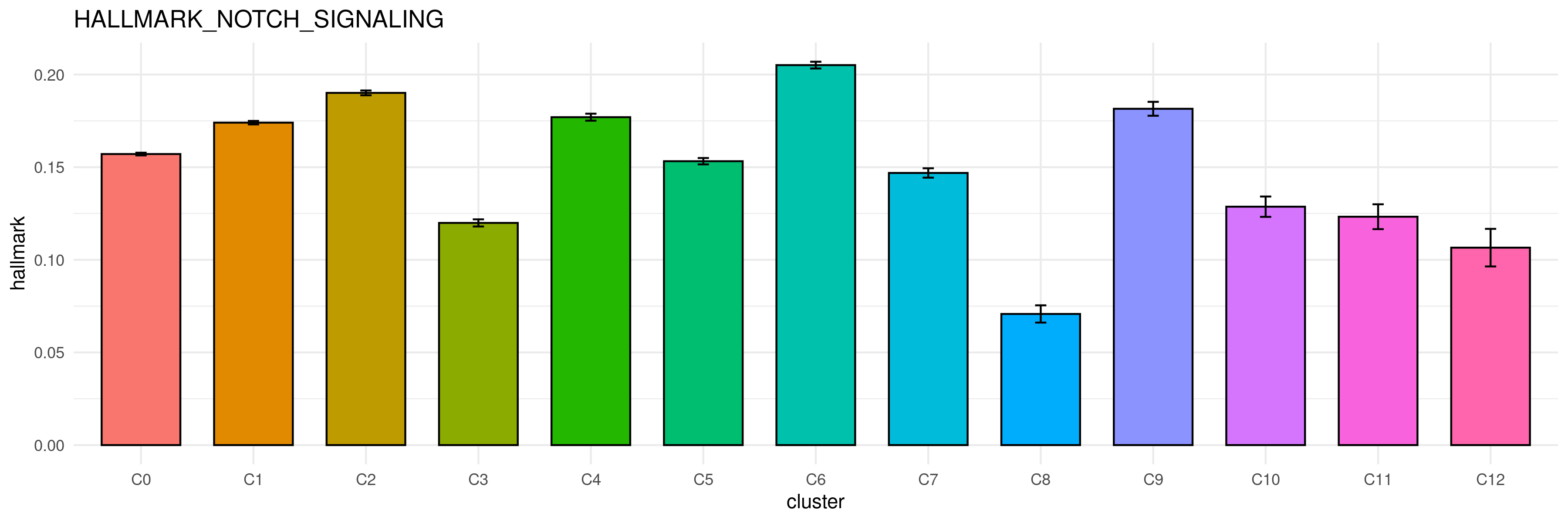

I

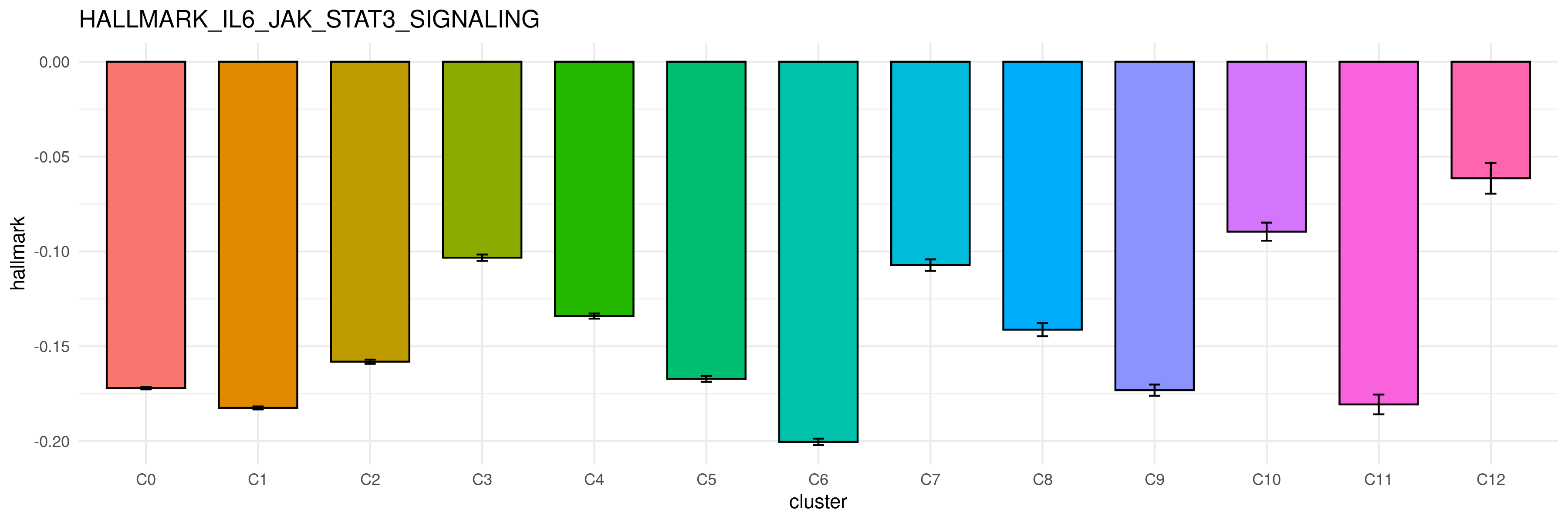

J

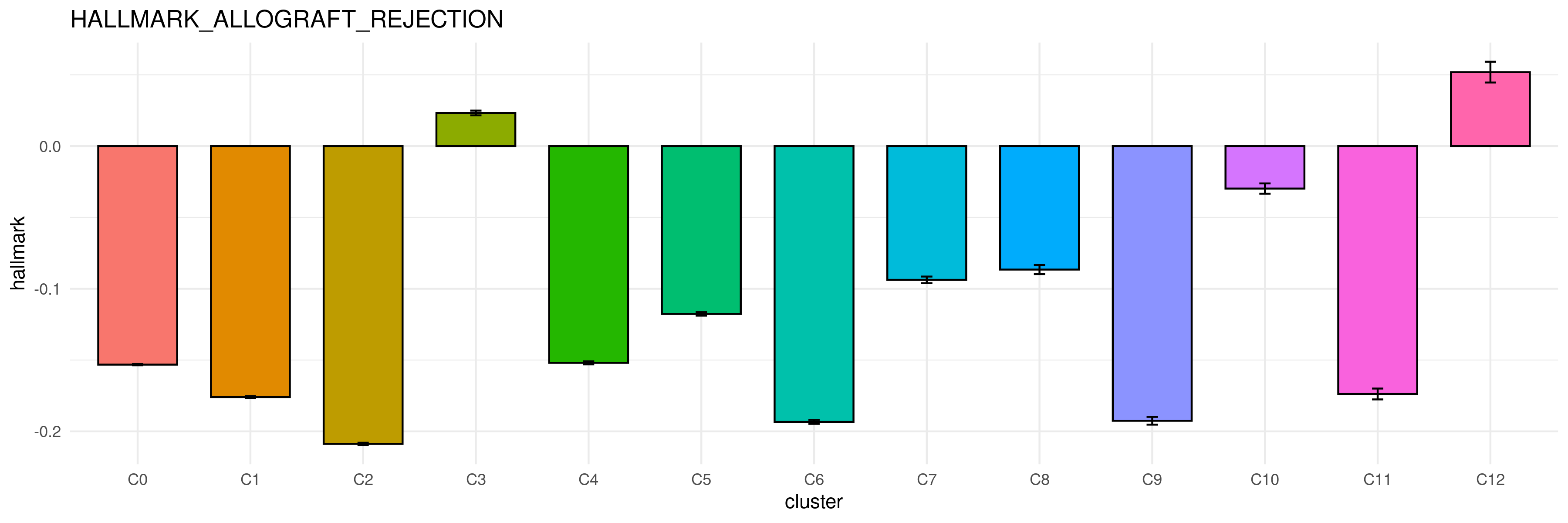

K

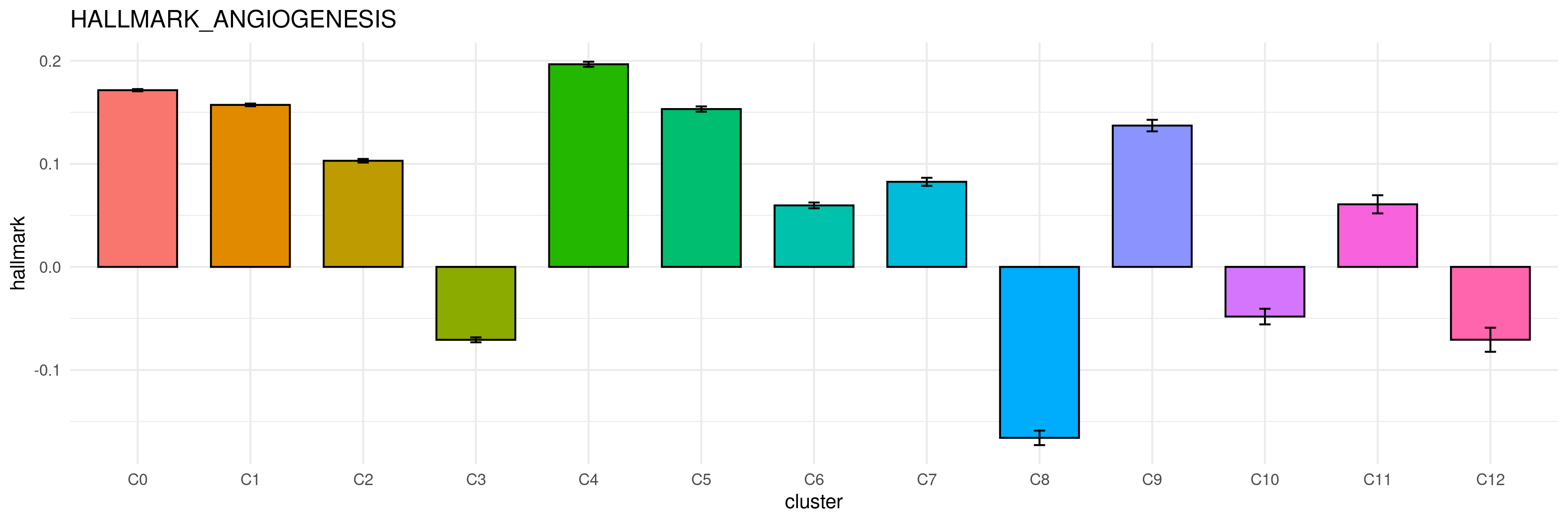

L

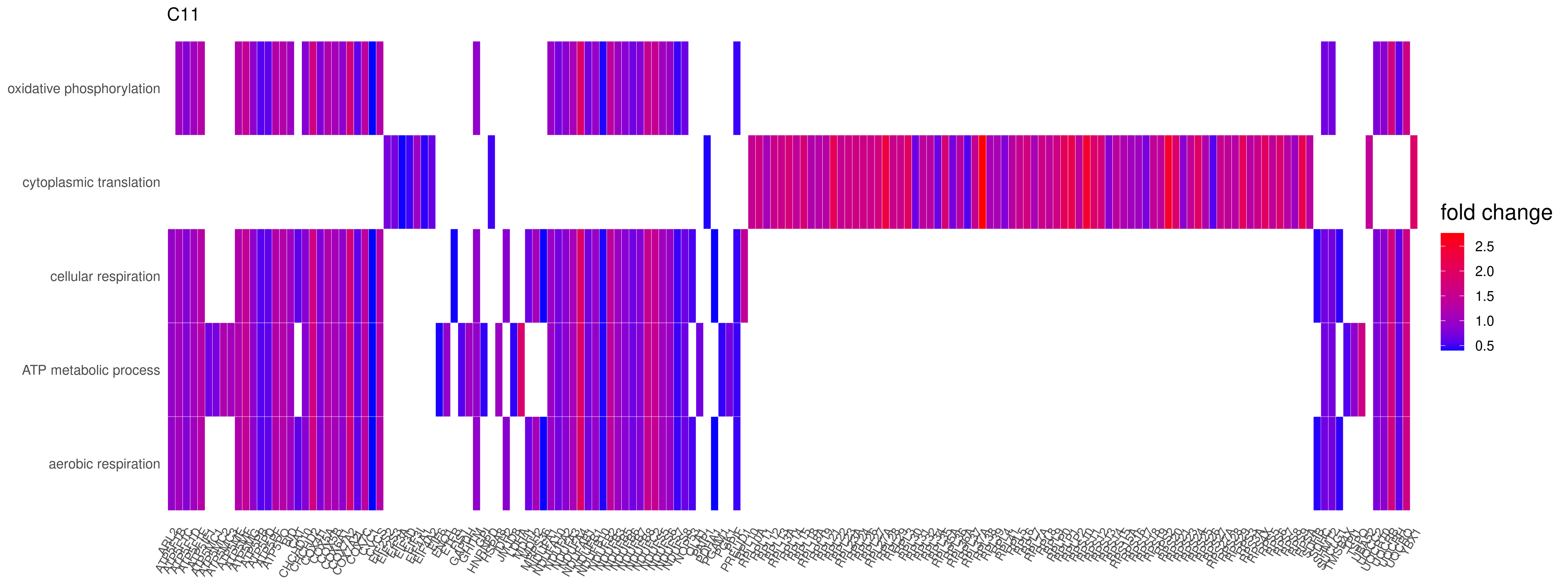

M

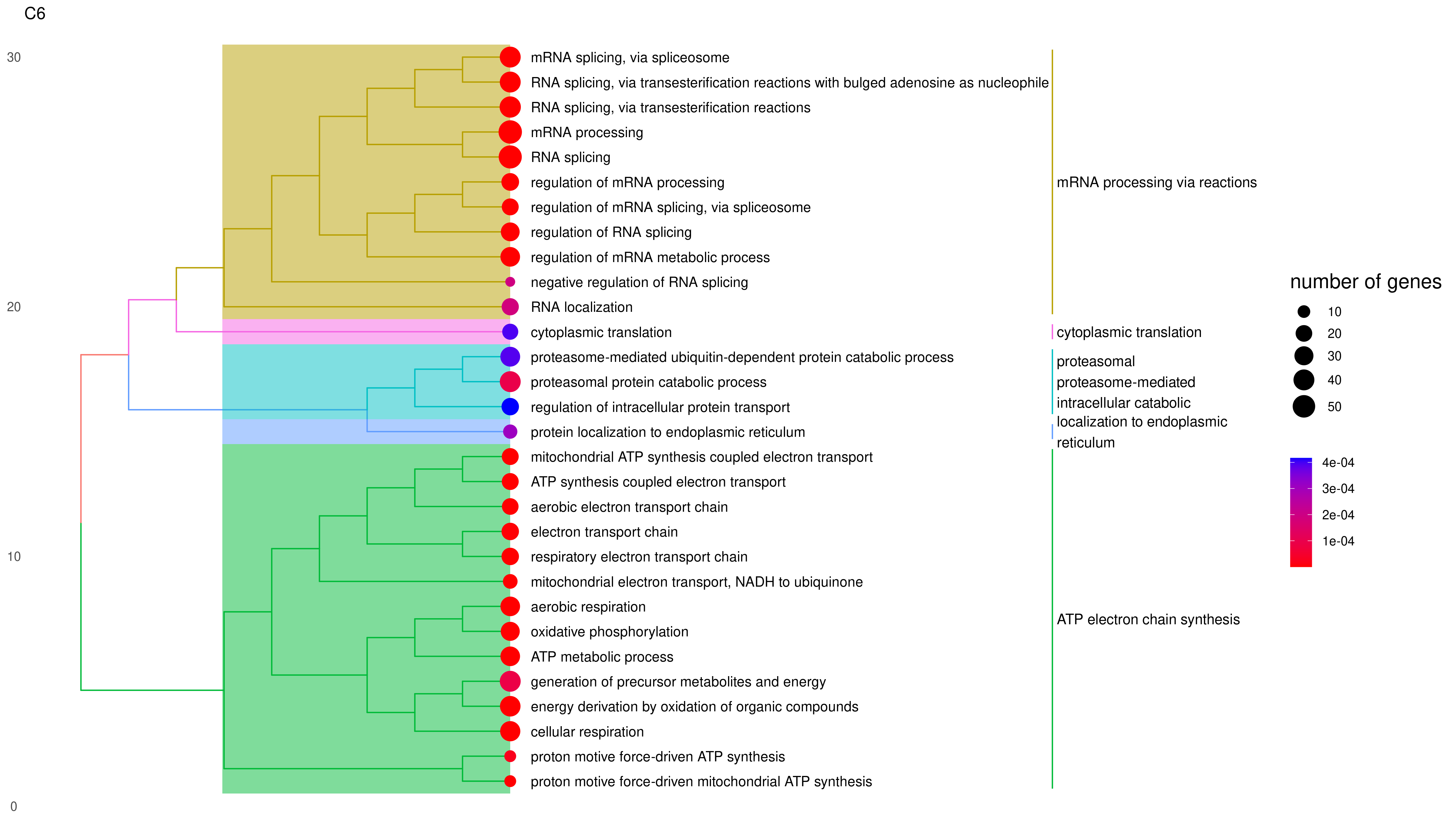

N

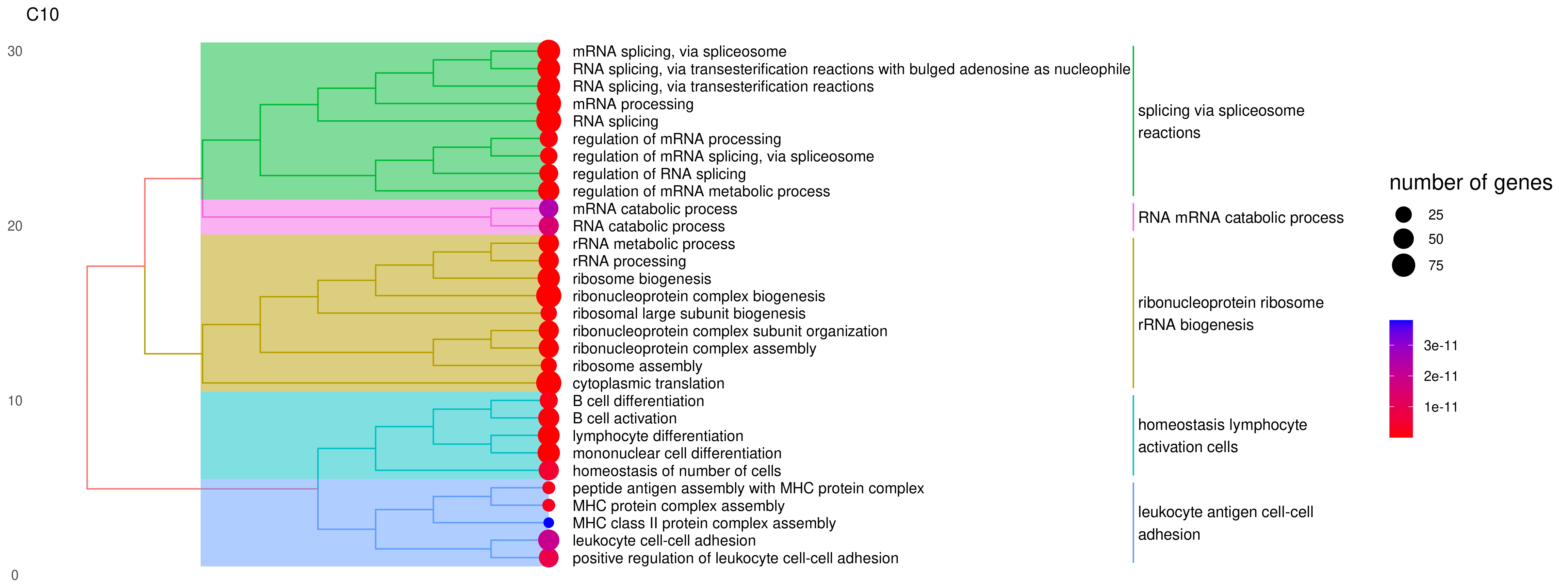

**Supplementary Figure 2**. Clustered platelets and their unique pathway expression changes.

(A - K) Bar plot of hallmark gene sets expression among clusters. A) MYC_targets_v1, B)MYC_targets_v1, C) Oxidative phosphorylation D) Glycolysis E) Inflammatory response F) Interferon alpha response, G) Interferon gamma response, H) Notch signaling I) IL6/JAK/STAT3 signaling. (L) Heatmap display fatal cluster C11 up-regulated genes and enriched pathways in gene ontology (GO). (M, N) Tree plot display survival cluster C6 and C10 enriched pathways in GO.

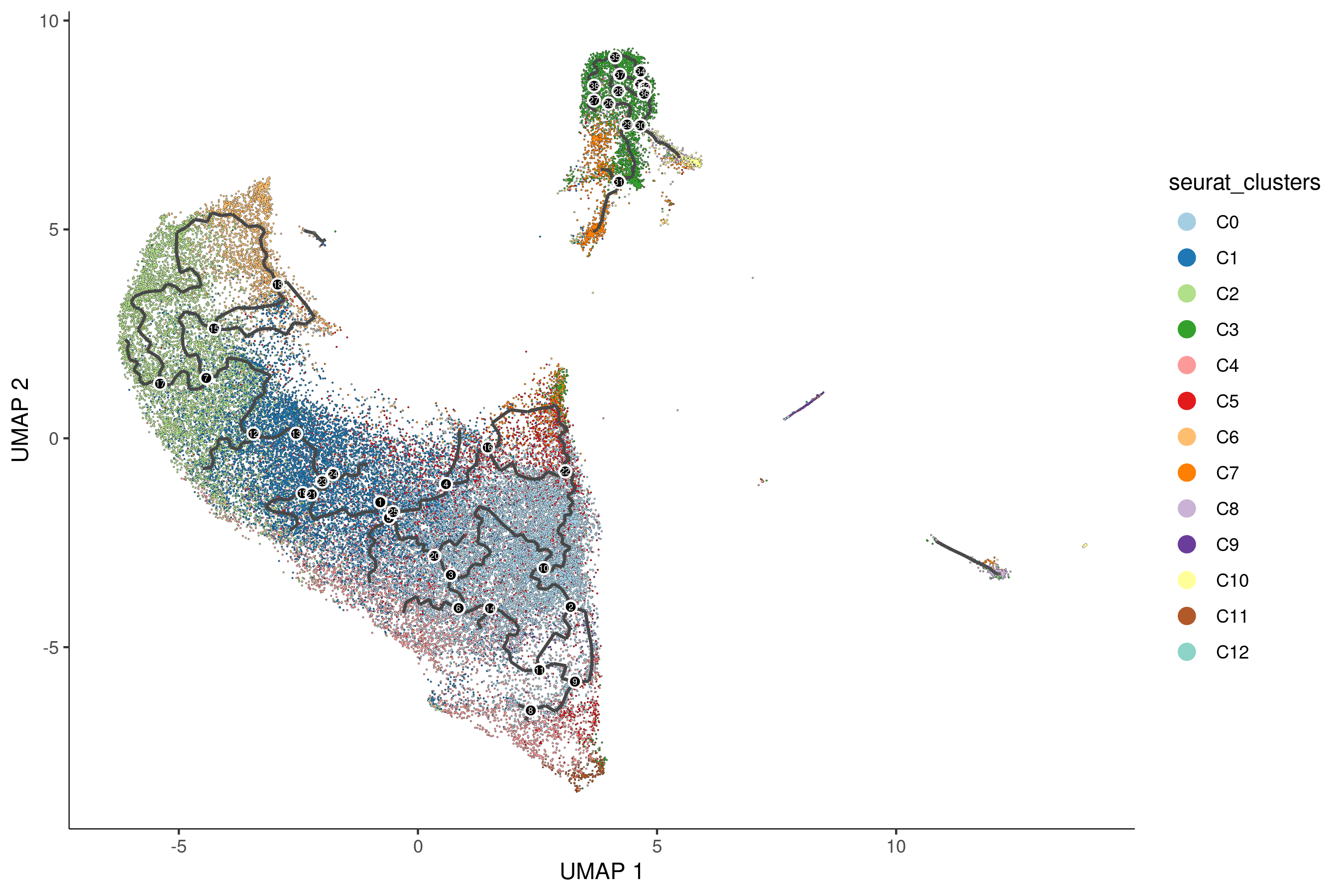

A

B

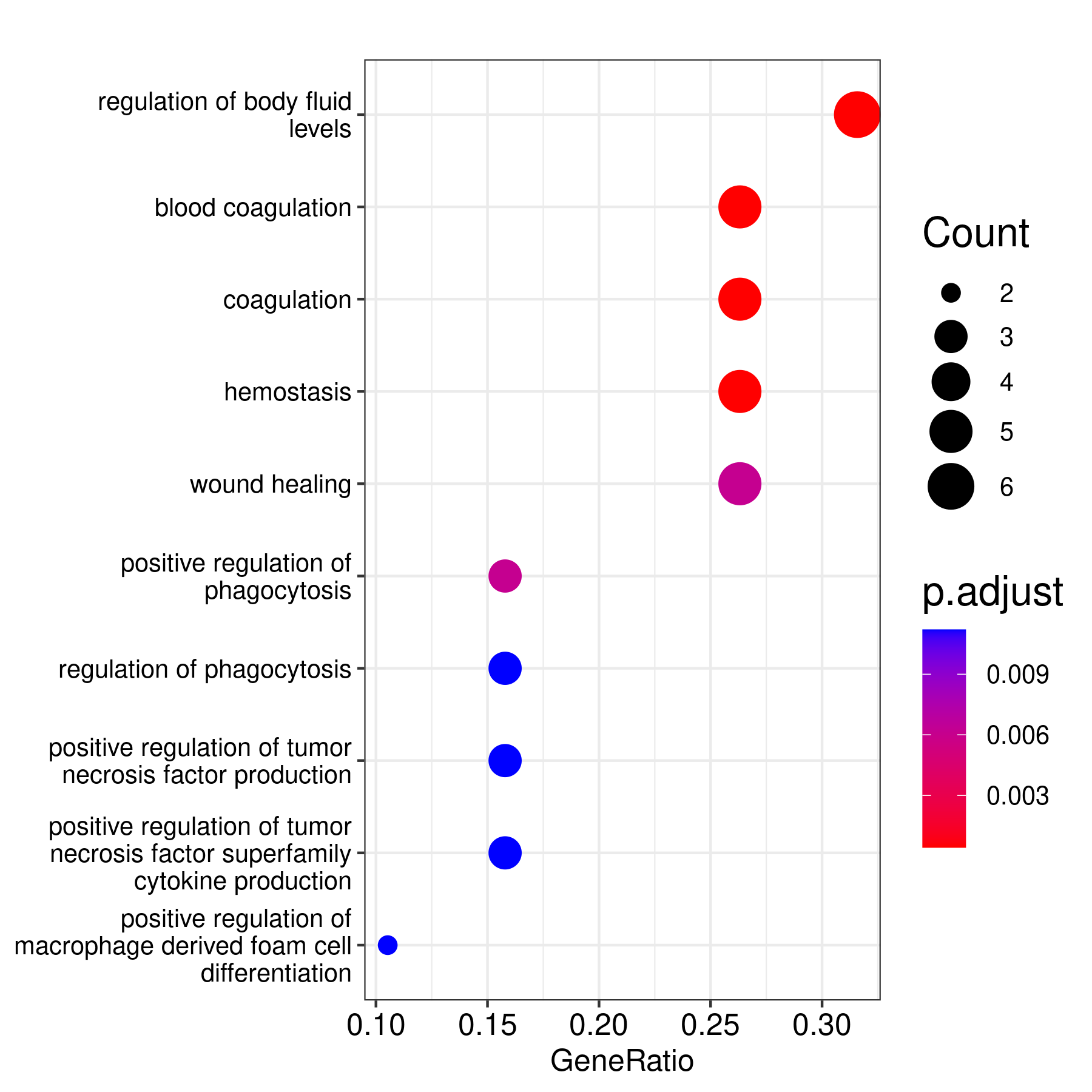

**Supplementary Figure 3**. (A) Pseudo-time plot of platelets from all clusters exhibiting trajectory fates. (B) The fatal gene module list enriched pathways from GO, KEGG, and Reactome databases. The circle size represents the gene counts included in the pathway; color represents the adjusted P value.

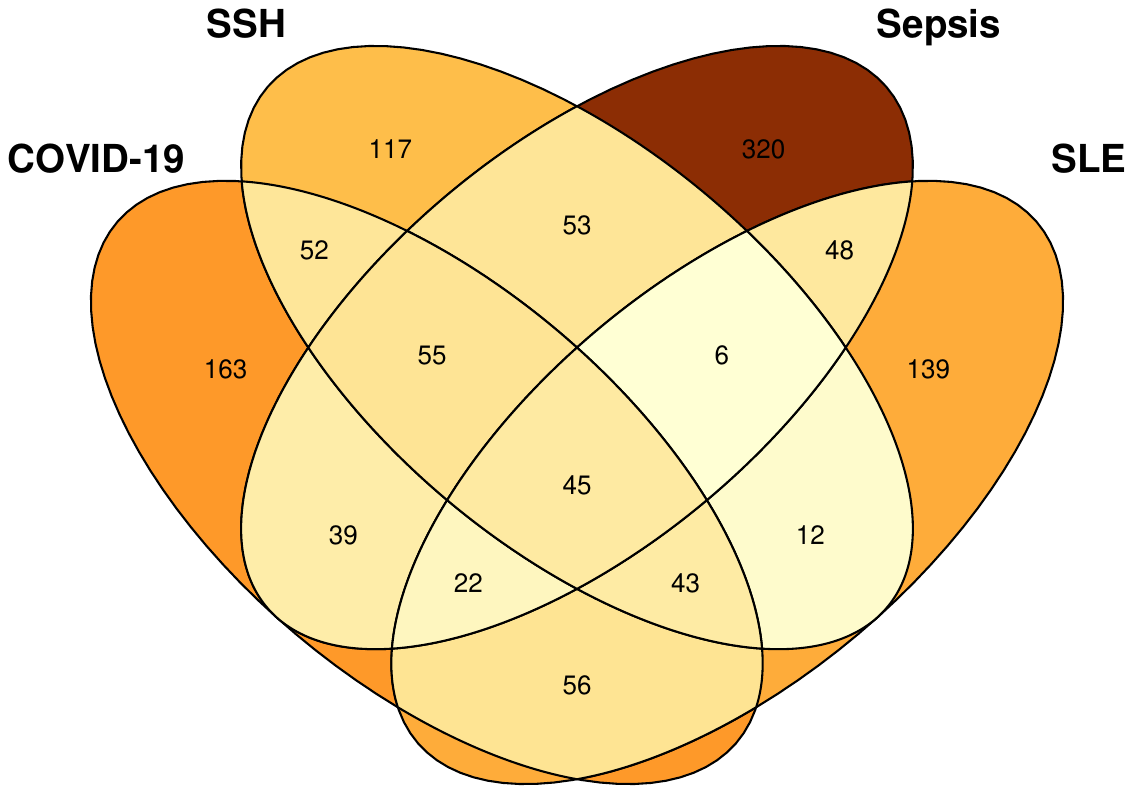

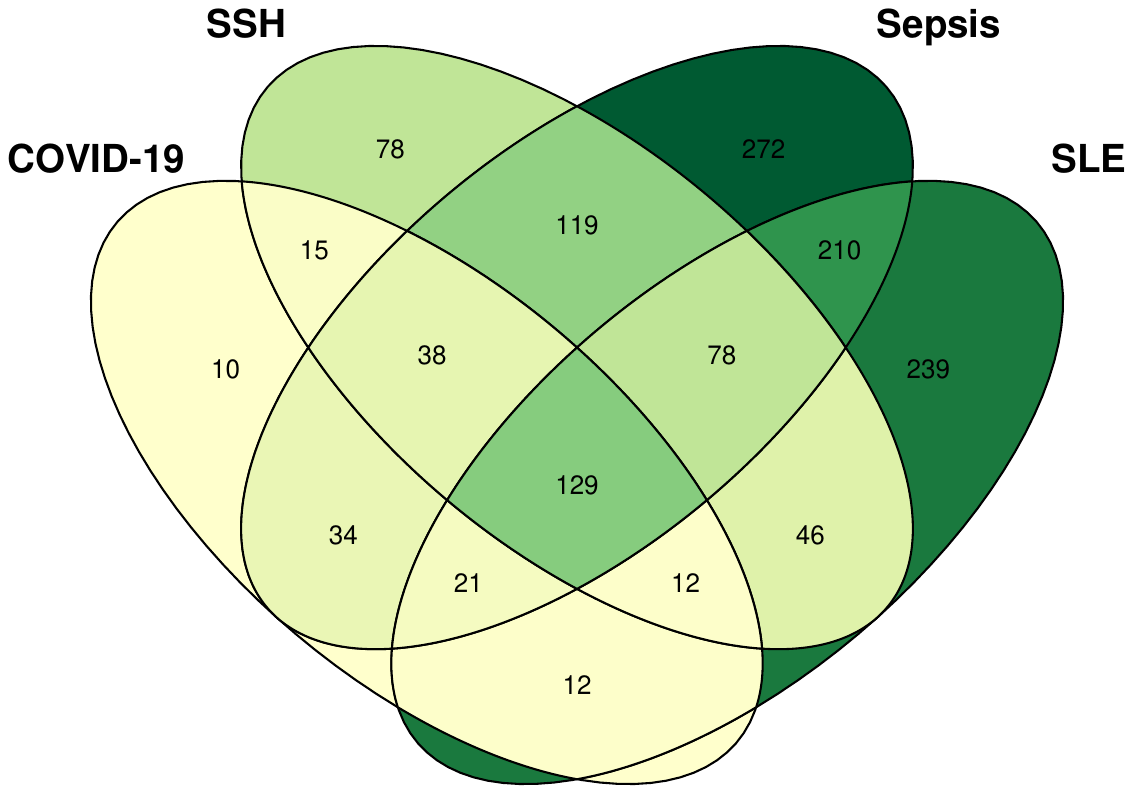

**Similar Symptoms Hospitalized**

**Similar Symptoms Hospitalized**

B

**Sepsis**

**SLE**

**Sepsis**

A

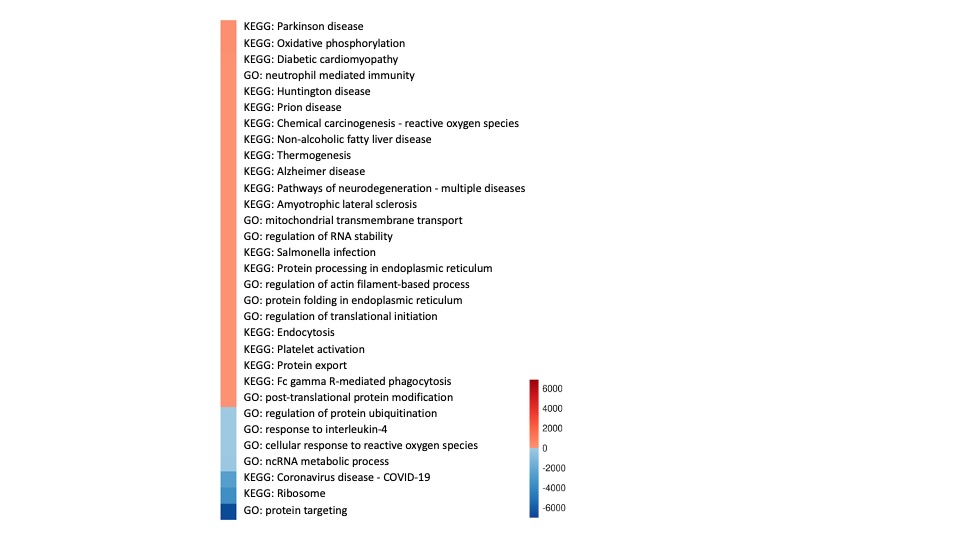

C

**Supplementary Figure 4**. **Platelets expression changes among healthy controls, sepsis, similar symptom hospitalized, COVID-19 and SLE patients**. Venn diagrams describing changes in differential expression genes (DEGs) from COVID-19, similar symptoms hospitalized (SSH), sepsis, and systemic sclerosis lupus (SLE) vs. healthy controls (HC), A) Up-regulated genes. B) Down-regulated genes. DEGs with adjusted p value below 0.05 and log2 fold change over 0.25 were used for generation of the VENN figure. (C) Heatmap illustration of COVID-19 and sepsis patients vs healthy contol pathways. Color are decided by the product of the COVID-19 and sepsis up/down-regulated enriched pathway log10 (adjusted p value). The GO terms were reduced to representative ones using Revigo [PMID: 21789182] (the cutoffs were similarity > 0.4) then overlapped the terms.

C

B

A

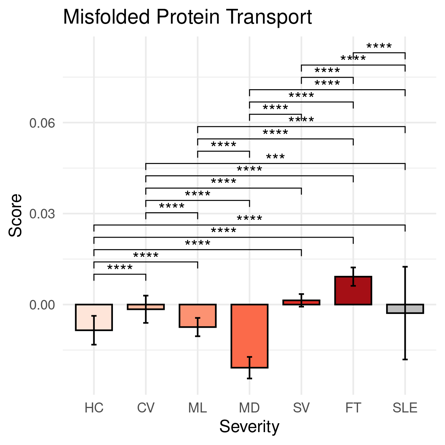

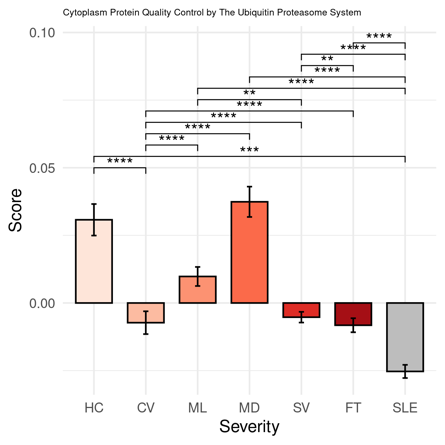

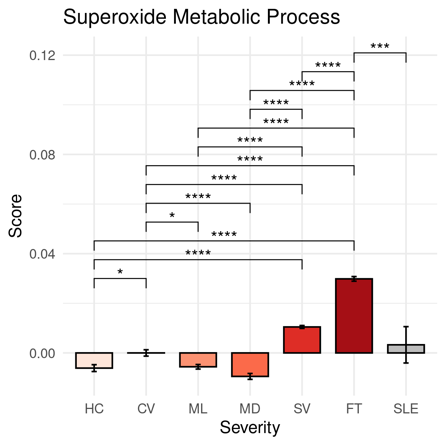

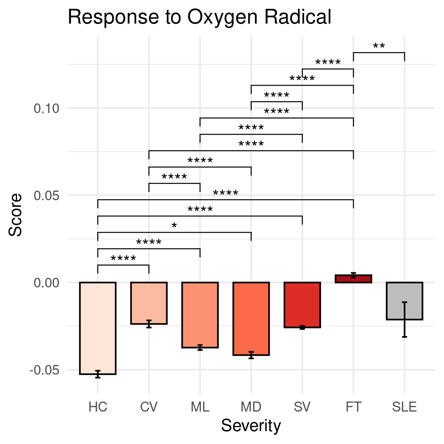

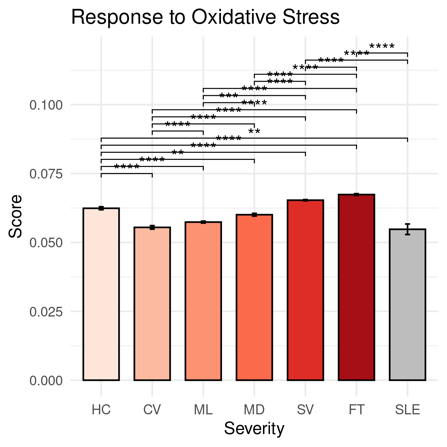

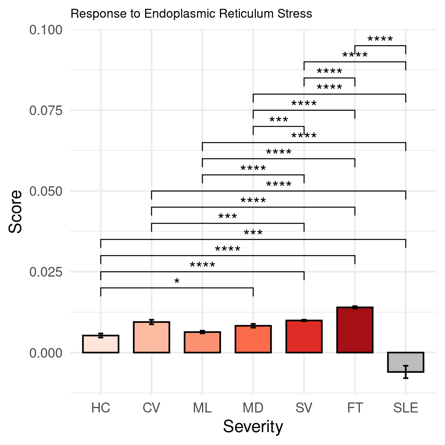

H

G

F

E

D

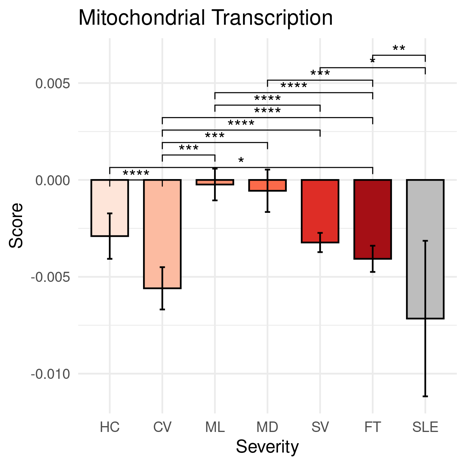

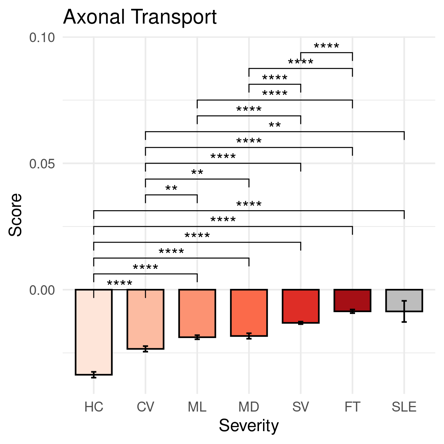

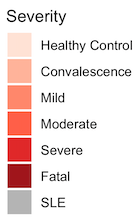

**Supplementary Figure 5**. **Platelet transcriptional changes under different disease severity related to neurodegeneration diseases**.

**(A– H)** Comparisons of pathway module scores across disease severities in platelets. The included modules contain genes related to **A)** Misfolded Protein Transport, **(B)** Cytoplasm Protein Quality Control by The Ubiquitin Proteasome System, **(C)** Superoxide Metabolic Process, **(D)** Response to Oxygen Radical, **(E)** Response to Oxidative Stress, (**F)** Response to Endoplasmic Reticulum Stress, **(G)** Mitochondrial Transcription**, (H)** Axonal Transport.

The differences in scores associated with adjusted *P*-values below 0.05, 0.01, 0.001, and 0.0001 are indicated as *, **, ***, and ****, respectively and not significant ones are not shown. The significance analysis was performed using Wilcoxon tests.

| Clusters | Severity | Number | Frequency |
| --- | --- | --- | --- |
| C0 | CV | 1107 | 0.082 |
| C0 | FT | 2876 | 0.214 |
| C0 | HC | 743 | 0.055 |
| C0 | MD | 1027 | 0.076 |
| C0 | ML | 1959 | 0.146 |
| C0 | SLE | 53 | 0.004 |
| C0 | SV | 5688 | 0.423 |
| C1 | CV | 879 | 0.083 |
| C1 | FT | 2101 | 0.2 |
| C1 | HC | 608 | 0.058 |
| C1 | MD | 949 | 0.09 |
| C1 | ML | 1839 | 0.175 |
| C1 | SLE | 51 | 0.005 |
| C1 | SV | 4102 | 0.39 |
| C2 | CV | 561 | 0.073 |
| C2 | FT | 1377 | 0.179 |
| C2 | HC | 305 | 0.04 |
| C2 | MD | 722 | 0.094 |
| C2 | ML | 1338 | 0.174 |
| C2 | SLE | 17 | 0.002 |
| C2 | SV | 3373 | 0.438 |
| C3 | CV | 81 | 0.021 |
| C3 | FT | 202 | 0.052 |
| C3 | HC | 636 | 0.165 |
| C3 | MD | 712 | 0.184 |
| C3 | ML | 962 | 0.249 |
| C3 | SLE | 4 | 0.001 |
| C3 | SV | 1267 | 0.328 |
| C4 | CV | 137 | 0.043 |
| C4 | FT | 1224 | 0.385 |
| C4 | HC | 139 | 0.044 |
| C4 | MD | 127 | 0.04 |
| C4 | ML | 264 | 0.083 |
| C4 | SLE | 4 | 0.001 |
| C4 | SV | 1284 | 0.404 |
| C5 | CV | 155 | 0.055 |
| C5 | FT | 520 | 0.185 |
| C5 | HC | 472 | 0.167 |
| C5 | MD | 310 | 0.11 |
| C5 | ML | 263 | 0.093 |
| C5 | SLE | 10 | 0.004 |
| C5 | SV | 1088 | 0.386 |
| C6 | CV | 168 | 0.072 |
| C6 | FT | 262 | 0.112 |
| C6 | HC | 18 | 0.008 |
| C6 | MD | 293 | 0.125 |
| C6 | ML | 375 | 0.16 |
| C6 | SLE | 11 | 0.005 |
| C6 | SV | 1211 | 0.518 |
| C7 | CV | 125 | 0.081 |
| C7 | FT | 211 | 0.137 |
| C7 | HC | 65 | 0.042 |
| C7 | MD | 150 | 0.097 |
| C7 | ML | 212 | 0.138 |
| C7 | SLE | 13 | 0.008 |
| C7 | SV | 764 | 0.496 |
| C8 | CV | 370 | 0.36 |
| C8 | FT | 86 | 0.084 |
| C8 | HC | 57 | 0.055 |
| C8 | MD | 15 | 0.015 |
| C8 | ML | 38 | 0.037 |
| C8 | SLE | 0 | 0 |
| C8 | SV | 462 | 0.449 |
| C9 | CV | 28 | 0.047 |
| C9 | FT | 263 | 0.442 |
| C9 | HC | 102 | 0.171 |
| C9 | MD | 9 | 0.015 |
| C9 | ML | 37 | 0.062 |
| C9 | SLE | 0 | 0 |
| C9 | SV | 156 | 0.262 |
| C10 | CV | 51 | 0.105 |
| C10 | FT | 62 | 0.127 |
| C10 | HC | 36 | 0.074 |
| C10 | MD | 8 | 0.016 |
| C10 | ML | 42 | 0.086 |
| C10 | SLE | 6 | 0.012 |
| C10 | SV | 282 | 0.579 |
| C11 | CV | 6 | 0.021 |
| C11 | FT | 221 | 0.784 |
| C11 | HC | 3 | 0.011 |
| C11 | MD | 0 | 0 |
| C11 | ML | 10 | 0.035 |
| C11 | SLE | 0 | 0 |
| C11 | SV | 42 | 0.149 |
| C12 | CV | 27 | 0.158 |
| C12 | FT | 9 | 0.053 |
| C12 | HC | 21 | 0.123 |
| C12 | MD | 8 | 0.047 |
| C12 | ML | 20 | 0.117 |
| C12 | SLE | 0 | 0 |
| C12 | SV | 86 | 0.503 |

**Supplementary Table 1**. Number and frequency of each cluster of platelets composed of each disease severity.

| Clusters | Outcomes | Number | Frequency |
| --- | --- | --- | --- |
| C0 | HC | 743 | 0.055 |
| C0 | FT | 2876 | 0.214 |
| C0 | S | 7135 | 0.53 |
| C0 | Unknown | 2699 | 0.201 |
| C1 | HC | 608 | 0.058 |
| C1 | FT | 2101 | 0.2 |
| C1 | S | 5527 | 0.525 |
| C1 | Unknown | 2293 | 0.218 |
| C2 | HC | 305 | 0.04 |
| C2 | FT | 1377 | 0.179 |
| C2 | S | 4325 | 0.562 |
| C2 | Unknown | 1686 | 0.219 |
| C3 | HC | 636 | 0.165 |
| C3 | FT | 202 | 0.052 |
| C3 | S | 2394 | 0.62 |
| C3 | Unknown | 632 | 0.164 |
| C4 | HC | 139 | 0.044 |
| C4 | FT | 1224 | 0.385 |
| C4 | S | 1106 | 0.348 |
| C4 | Unknown | 710 | 0.223 |
| C5 | HC | 472 | 0.167 |
| C5 | FT | 520 | 0.185 |
| C5 | S | 1190 | 0.422 |
| C5 | Unknown | 636 | 0.226 |
| C6 | HC | 18 | 0.008 |
| C6 | FT | 262 | 0.112 |
| C6 | S | 1675 | 0.716 |
| C6 | Unknown | 383 | 0.164 |
| C7 | HC | 65 | 0.042 |
| C7 | FT | 211 | 0.137 |
| C7 | S | 1020 | 0.662 |
| C7 | Unknown | 244 | 0.158 |
| C8 | HC | 57 | 0.055 |
| C8 | FT | 86 | 0.084 |
| C8 | S | 774 | 0.753 |
| C8 | Unknown | 111 | 0.108 |
| C9 | HC | 102 | 0.171 |
| C9 | FT | 263 | 0.442 |
| C9 | S | 156 | 0.262 |
| C9 | Unknown | 74 | 0.124 |
| C10 | HC | 36 | 0.074 |
| C10 | FT | 62 | 0.127 |
| C10 | S | 350 | 0.719 |
| C10 | Unknown | 39 | 0.08 |
| C11 | HC | 3 | 0.011 |
| C11 | FT | 221 | 0.784 |
| C11 | S | 45 | 0.16 |
| C11 | Unknown | 13 | 0.046 |
| C12 | HC | 21 | 0.123 |
| C12 | FT | 9 | 0.053 |
| C12 | S | 53 | 0.31 |
| C12 | Unknown | 88 | 0.515 |

**Supplementary Table 2**. Number and frequency of each cluster of platelets composed of each outcome situation.

| Cluster | Gene | avg_log2FC | pct.1 | pct.2 | p_val | p_val_adj |
| --- | --- | --- | --- | --- | --- | --- |
| C0 | MYL9 | 1.44979842 | 0.927 | 0.676 | 0 | 0 |
| C0 | FCER1G | 1.3785031 | 0.695 | 0.431 | 0 | 0 |
| C0 | PARVB | 1.36861383 | 0.659 | 0.433 | 0 | 0 |
| C1 | TMEM140 | 1.08200129 | 0.626 | 0.378 | 0 | 0 |
| C1 | PNMA1 | 1.05784379 | 0.356 | 0.22 | 0 | 0 |
| C1 | CDKN1A | 1.04413495 | 0.608 | 0.41 | 0 | 0 |
| C2 | DEPP1 | 2.46460805 | 0.233 | 0.014 | 0 | 0 |
| C2 | INKA1 | 2.3932049 | 0.64 | 0.103 | 0 | 0 |
| C2 | FAM110A | 2.21126078 | 0.805 | 0.225 | 0 | 0 |
| C3 | JUNB | 3.28521179 | 0.807 | 0.137 | 0 | 0 |
| C3 | JUN | 3.27880346 | 0.79 | 0.12 | 0 | 0 |
| C3 | RPL34 | 3.17377804 | 0.975 | 0.319 | 0 | 0 |
| C4 | HPSE | 1.46464595 | 0.6 | 0.148 | 0 | 0 |
| C4 | WFDC1 | 1.41410805 | 0.354 | 0.058 | 0 | 0 |
| C4 | PF4 | 1.31987935 | 0.998 | 0.93 | 0 | 0 |
| C5 | JCHAIN | 1.46558029 | 0.258 | 0.105 | 7.35E-164 | 9.60E-160 |
| C5 | TMSB10 | 0.31795166 | 0.651 | 0.491 | 7.48E-66 | 9.77E-62 |
| C5 | PSME2 | 0.28101928 | 0.076 | 0.104 | 1.21E-64 | 1.58E-60 |
| C6 | CRBN | 2.57584885 | 0.717 | 0.227 | 0 | 0 |
| C6 | CD58 | 2.54909621 | 0.334 | 0.049 | 0 | 0 |
| C6 | SRSF3 | 2.50170105 | 0.639 | 0.197 | 0 | 0 |
| C7 | S100A8 | 2.77269395 | 0.73 | 0.434 | 0 | 0 |
| C7 | MALAT1 | 2.75406188 | 0.965 | 0.655 | 0 | 0 |
| C7 | NEAT1 | 2.34567652 | 0.645 | 0.245 | 0 | 0 |
| C8 | RPLP0 | 3.02681342 | 0.967 | 0.261 | 0 | 0 |
| C8 | RPS6 | 2.84360105 | 0.974 | 0.266 | 0 | 0 |
| C8 | RPS23 | 2.84118611 | 0.994 | 0.323 | 0 | 0 |
| C9 | HBB | 6.9482949 | 0.576 | 0.212 | 0 | 0 |
| C9 | HBA2 | 6.53307512 | 0.45 | 0.084 | 0 | 0 |
| C9 | HBA1 | 5.67655265 | 0.501 | 0.101 | 0 | 0 |
| C10 | CD74 | 3.89201311 | 0.986 | 0.299 | 0 | 0 |
| C10 | HLA-DRA | 3.41506579 | 0.899 | 0.094 | 0 | 0 |
| C10 | CD79A | 2.77837866 | 0.674 | 0.019 | 0 | 0 |
| C11 | TPT1 | 2.24007978 | 0.996 | 0.635 | 1.13E-210 | 1.48E-206 |
| C11 | POLR2L | 2.34141381 | 0.904 | 0.143 | 7.60E-195 | 9.93E-191 |
| C11 | CSRP1 | 1.94266214 | 0.798 | 0.093 | 3.92E-171 | 5.12E-167 |
| C12 | MALAT1 | 2.74811184 | 1 | 0.663 | 1.09E-122 | 1.42E-118 |
| C12 | RPL3 | 2.48307029 | 0.971 | 0.29 | 1.54E-117 | 2.02E-113 |
| C12 | RPS16 | 2.21622308 | 0.977 | 0.26 | 7.54E-106 | 9.85E-102 |

**Supplementary Table 3**. Up-regulated genes in each platelet cluster compared to other clusters of platelets.

| COVID-19, SSH, sepsis, SLE compare to HC | |
| --- | --- |
| Overlapped up-regulated genes | Overlapped down-regulated genes |
| AP2B1 | ANXA1 |
| ATP5F1E | BTG1 |
| ATP5MPL | BTG2 |
| BEX3 | CABP5 |
| BRK1 | CD3E |
| C9orf16 | CD48 |
| CASP4 | CD52 |
| CD226 | CD7 |
| CYTOR | COMMD6 |
| DNAJC15 | CORO1A |
| DSTN | CRIP1 |
| DYNLRB1 | CXCR4 |
| EIF1 | DDX5 |
| FCER1G | DUSP1 |
| GLA | EEF1A1 |
| GNG5 | EEF1B2 |
| GPX4 | EEF1D |
| GTF2A2 | EEF2 |
| HSPB1 | EIF2S3 |
| IFI27L2 | EIF3E |
| IFITM2 | EIF3F |
| IFITM3 | EIF3H |
| ISCA1 | EIF3K |
| KIFAP3 | EIF3L |
| LCN2 | ETS1 |
| MAPRE1 | FAU |
| NAA38 | FOS |
| NAP1L1 | GIMAP7 |
| NDUFS5 | GRHL1 |
| OST4 | GTPBP2 |
| RHEB | HNRNPA1 |
| RHOC | HNRNPDL |
| S100A8 | HSPA8 |
| S100A9 | ICAM3 |
| SERF2 | IDS |
| SMIM27 | IER2 |
| STMP1 | IL32 |
| TBC1D15 | IL7R |
| TIMP1 | JUN |
| TMEM219 | JUNB |
| UBL5 | KLF2 |
| UQCR11 | LCK |
| XK | LDHB |
| YWHAE | LEF1 |
| ZFAND3 | LEPROTL1 |
|  | LIMD2 |
|  | LINC00861 |
|  | LITAF |
|  | LSP1 |
|  | LTB |
|  | MAL |
|  | MALAT1 |
|  | MRFAP1 |
|  | NACA |
|  | NOSIP |
|  | NPM1 |
|  | ODC1 |
|  | PABPC1 |
|  | PFDN5 |
|  | PTPRC |
|  | RACK1 |
|  | RGCC |
|  | RPL10 |
|  | RPL10A |
|  | RPL11 |
|  | RPL12 |
|  | RPL13 |
|  | RPL14 |
|  | RPL17 |
|  | RPL18 |
|  | RPL18A |
|  | RPL19 |
|  | RPL22 |
|  | RPL23A |
|  | RPL24 |
|  | RPL26 |
|  | RPL28 |
|  | RPL29 |
|  | RPL3 |
|  | RPL30 |
|  | RPL32 |
|  | RPL34 |
|  | RPL35 |
|  | RPL36 |
|  | RPL37 |
|  | RPL39 |
|  | RPL4 |
|  | RPL5 |
|  | RPL6 |
|  | RPL7 |
|  | RPL7A |
|  | RPL8 |
|  | RPLP0 |
|  | RPLP2 |
|  | RPS12 |
|  | RPS13 |
|  | RPS14 |
|  | RPS15 |
|  | RPS15A |
|  | RPS16 |
|  | RPS18 |
|  | RPS19 |
|  | RPS2 |
|  | RPS21 |
|  | RPS23 |
|  | RPS24 |
|  | RPS25 |
|  | RPS26 |
|  | RPS27A |
|  | RPS28 |
|  | RPS3 |
|  | RPS3A |
|  | RPS4X |
|  | RPS5 |
|  | RPS6 |
|  | RPS7 |
|  | RPS8 |
|  | RPSA |
|  | SARAF |
|  | SLC25A6 |
|  | SNRPD2 |
|  | SSR2 |
|  | TCF7 |
|  | TNS1 |
|  | TOMM20 |
|  | TOMM7 |
|  | TSPAN18 |
|  | TUBA1C |
|  | ZFP36L2 |

**Supplementary Table 4**. Consistently up/down regulated genes in COVID-19, similar symptom hospitalized (SSH), sepsis, and SLE patients compared to healthy controls (HC).
